# The co-localizing Zorya II, Druantia III, and ARMADA II defense systems on O-island 172 confer synergistic anti-phage defense in enterohemorrhagic *Escherichia coli*

**DOI:** 10.1101/2025.09.22.677730

**Authors:** Huihui Li, Vu Van Loi, Costanza Borelli, Stephanie Himpich, Michaela Projahn, Lena M. Grass, Thi Phuong Thao Nguyen, Markus C. Wahl, Haike Antelmann

**Affiliations:** Freie Universität Berlin, Institute of Biology-Microbiology, D-14195 Berlin, Germany; Freie Universität Berlin, Laboratory of Structural Biochemistry, D-14195 Berlin, Germany; National Reference Laboratory for Escherichia coli Including VTEC, German Federal Institute for Risk Assessment, 12277 Berlin, Germany; Helmholtz-Zentrum Berlin für Materialien und Energie, Macromolecular Crystallography, D-12489 Berlin, Germany

**Keywords:** Enterohemorrhagic *Escherichia coli* O157:H7, EDL933, OI-172, anti-phage defense systems, Zorya II, Druantia III, ARMADA II, synergy

## Abstract

In this study, we show that the enterohemorrhagic *Escherichia coli* (EHEC) strain EDL933 Δ*stx1/2* is highly resistant to phages. The genes encoding the three phage defense systems Zorya II, Druantia III, and ARMADA II were found to co-localize on O-island 172 (OI-172) in EDL933. The three phage defense systems co-occur in 426 *E. coli* strains comprising 66 distinct serotypes, including 277 O157:H7 isolates and 117 diverse non-pathogenic strains. Efficiency of plating (EOP) assays were used to explore the phage spectrum and synergy of the three defense systems and their contribution to the phage resistance of strain EDL933 Δ*stx1/2*. Using Δ*zorABE,* Δ*druHE*, and Δ*armABCD* single system deletion mutants in EDL933 Δ*stx1/2*, we showed that Druantia III and ARMADA II protect to different extents against a broad spectrum of phages, including the *Drexlerviridae*, *Siphoviridae*, *Demerecviridae*, *Vequintavirinae* and *Autographiviridae*, but not against hypermodified *Tevenvirinae*. Additionally, Zorya II caused strong protection only against *Autographiviridae*. Furthermore, EOP assays of combined system mutants revealed strong synergistic interactions of Druantia III and ARMADA II to provide more robust immunity against a similar phage spectrum than the additive protection of the individual systems. In contrast, Druantia III and Zorya II act only weakly synergistically against a few phages. Altogether, our results revealed that Druantia III and ARMADA II are responsible for most of the phage resistance of EDL933 Δ*stx1/2*, whereas Zorya II provides additional immunity against podoviruses. Future studies are underway to elucidate the molecular basis of the synergistic interactions between the Druantia III and ARMADA II defense systems.

**IMPORTANCE:** Shiga toxin-producing *E. coli* (STEC) O157:H7 strains cause life-threatening diseases, such as hemorrhagic colitis and the hemolytic uremic syndrome. Currently, EHEC infections can be only treated symptomatically, since antibiotics are not recommended due to induction of the Shiga toxin. While phage therapy could offer a treatment option, we show that the EHEC strain EDL933 Δ*stx1/2* is highly resistant to many phages. EOP assays of defense mutants in the host strain revealed that the co-occurring defense systems Zorya II, Druantia III, and ARMADA II on OI-172 confer most of the phage resistance in EHEC. Moreover, our data uncovered that Druantia III and ARMADA II act synergistically in the anti-phage defense in EDL933 Δ*stx1/2*, explaining the robust immunity against a broad spectrum of phages. These results support the idea that the design of novel inhibitors against the Druantia III and ARMADA II systems could be combined with phage therapies to efficiently eradicate EHEC.

## INTRODUCTION

Enterohemorrhagic *Escherichia coli* (EHEC) O157:H7 are foodborne and zoonotic pathogens that are transmitted to humans through contaminated milk, meat, or plant-based food and may cause gastrointestinal infections, ranging from diarrhoea to life-threatening hemorrhagic colitis and the hemolytic uremic syndrome (HUS) (1–5). EHEC strains are a subgroup of Shiga toxin-producing *E. coli* (STEC), which cause severe outbreaks worldwide at a low infectious dose (<100 cfu) with many hospitalizations and a death rate of 3-5% (3, 6–9). The Robert Koch Institute (RKI) institute in Berlin has recently reported an EHEC/HUS outbreak in Mecklenburg-Vorpommern during August and September of >107 EHEC infections, including at least 53 patients infected with the rare O45:H2 serotype https://www.rki.de/DE/Home/home_node.html.

However, treatment options for EHEC infections with antibiotics are very limited, since many antibiotics induce the Shiga toxin as the major virulence factor, which enhances the risk for HUS development (10, 11). In addition, current therapies with coliphages against O157:H7 infections only led to modest bacterial killing, indicating that more specific phages against EHEC have to be isolated and alternative treatment strategies developed (12). Thus far, >130 anti-EHEC phages have been isolated from faecal samples of infected human patients and cattle, contaminated waste water, food, and sewage (13, 14). Many anti-EHEC phages had lytic activities against the O157:H7 serotype and other Gram-negative pathogens, e.g. *Shigella* and *Salmonella*, indicating a broad host range and making them ideal candidates for application in the food industry and medicine (13–16).

However, the constant arms race between bacteria and their phages has driven the evolution of hundreds of anti-phage defense systems in bacteria that provide innate and adaptive immunity towards phage infection at various stages of the viral infection cycle (17–21). The mechanisms of prokaryotic phage defense systems are remarkably complex, including sensing of phage DNA or proteins, signaling through production of small molecules, depletion of nucleotides, chemical defense or systems sensing the integrity of the host cell upon phage infection (17–21). The most abundant anti-phage defenses in bacteria are nuclease-based systems involved in degradation of phage nucleic acids, including restriction/modification (RM) systems, the Defense Island System Associated with the RM (DISARM), CRISPR-Cas, Zorya and Druantia defense systems (17–21). In the type I-III RM systems, methyltransferases (MTases) methylate adenine or cytosine bases in the host genome to discriminate non-methylated phage DNA, which is degraded by cognate restriction endonucleases (REs) at the unmodified restriction sites (17–19).

In the DISARM system, unmodified DNA substrates with single-stranded overhangs are recognized by the DrmAB helicase complex (22). However, no nuclease activity has so far been observed for DrmAB or the PLD-containing protein DrmC, and DrmC was not essential for DISARM anti-phage defense in vivo (22–24). The Druantia anti-phage defense systems were classified into Druantia types I, II, and III, which share the large DExH box DNA helicase/nuclease DruE (25), belonging to the YprA helicase family (26). However, the three Druantia subtypes have different accessory proteins, such as DruH in Druantia III (25).

Zorya defense systems are classified into Zorya types I, II, and III, which include a conserved ion-driven ZorA_5_B_2_ core complex containing the peptidoglycan-binding ZorB subunits and an elongated cytoplasmic tail assembled by the five ZorA subunits. The complex rotates and recruits different effector components for phage DNA degradation, including ZorC and the predicted DEAD box helicase ZorD (Zorya I), the HNH endonuclease ZorE (Zorya II), and the DUF3348 and DUF2894 domain proteins ZorF and ZorG (Zorya III) (21, 25, 27, 28). In Zorya II, the N-terminal helix of ZorB and the pentameric ZorA tail are shorter compared to Zorya I (28). The effector of the Zorya II system was identified as the ZorE nickase, which targets phage and chromosomal DNA (28).

Moreover, prokaryotic immune systems were shown to often co-localize in so-called defense islands due to the integration of mobile genetic elements, such as phages, in distinct genomic hotspots (17, 18, 20, 25, 29, 30). The co-occurrence of phage defense systems in *E. coli* has been associated with their synergistic anti-phage activities, as demonstrated for the Zorya II and Druantia III defense systems (31). Synergistic immunity has been shown to provide an evolutionary advantage against host-specific phages at the population level (31).

Recent studies in pathogenic antibiotic-resistant *Pseudomonas aeruginosa* isolates revealed a correlation between the number of phage defense systems with their levels of phage resistance, indicating additive and synergistic interactions in the specificity against the targeted phage spectrum (32, 33).

In this work, we found that the EHEC strain EDL933 Δs*tx1/2* exhibits strong resistance towards many phages of the BASEL phage collection (34), which efficiently lyse *E. coli* K-12 strains. DefenseFinder predicted three phage defense systems, which are encoded on O-island 172 (OI-172), including Zorya II, Druantia III, and a recently described ARMADA II system (DISARM-related Antiviral defense array) (26). This OI-172-associated phage defense island is distributed in O157:H7 strains and several non-pathogenic *E. coli* of the NCBI RefSeq collection. Using efficiency of plating (EOP) and growth assays of single and combined defense system mutants, we analyzed the phage spectra and the synergy of the three defense systems, revealing additive and synergistic interactions of Druantia III and ARMADA II compared to each system alone. Overall, our results identified a hitherto uncharacterized defense system in EHEC, ARMADA II, and demonstrated that the phage defense island on OI-172 is responsible for the strong phage resistance of EDL933 Δs*tx1/2*.

## RESULTS

### Identification of a phage defense island on OI-172 in EHEC O157:H7 strain EDL933

Using DefenseFinder (20), three phage defense systems were predicted to be encoded at the nine-gene locus *z5894-z5902* on OI-172 of *E. coli* O157:H7 strain EDL933, including Zorya II (ZorA, ZorB, and ZorE), Druantia III (DruH, DruE), and a DISARM I-related system, named as ARMADA II (ArmA, ArmB, ArmC, and ArmD) **(Fig. 1; Table S1)** (25). The Druantia III system encompasses the unknown function protein DruH (Z5897) and the very large ATP-dependent DExH box helicase/nuclease DruE (Z5898) containing DUF1998 and two PLD-like domains in the C-terminal region, which was recently classified as YprA family helicase (26) **(Fig. 1; Table S1)**. The tandem PLD-like domains are replaced by a single non-PLD endonuclease-like domain in Druantia I and II systems (25). We note that DruE proteins, therefore, resemble fusions of the DISARM I components DrmA (helicase), DrmB (DUF1998-containing protein), and either duplicated DrmC (PLD-containing protein) units (Druantia III) or single non-PLD endonuclease-like domains (Druantia I and II). Druantia III is widely distributed in pathogenic proteobacteria, including EHEC O157:H7 and O145:H28 strains, *P. aeruginosa, Salmonella paratyphi*, *Vibrio* sp., *Klebsiella pneumoniae*, and *Acinetobacter baumannii* (25, 35). Using EOP assays, Druantia III was shown to provide a broad range of defense against >70 phages (35).

**Fig 1.**
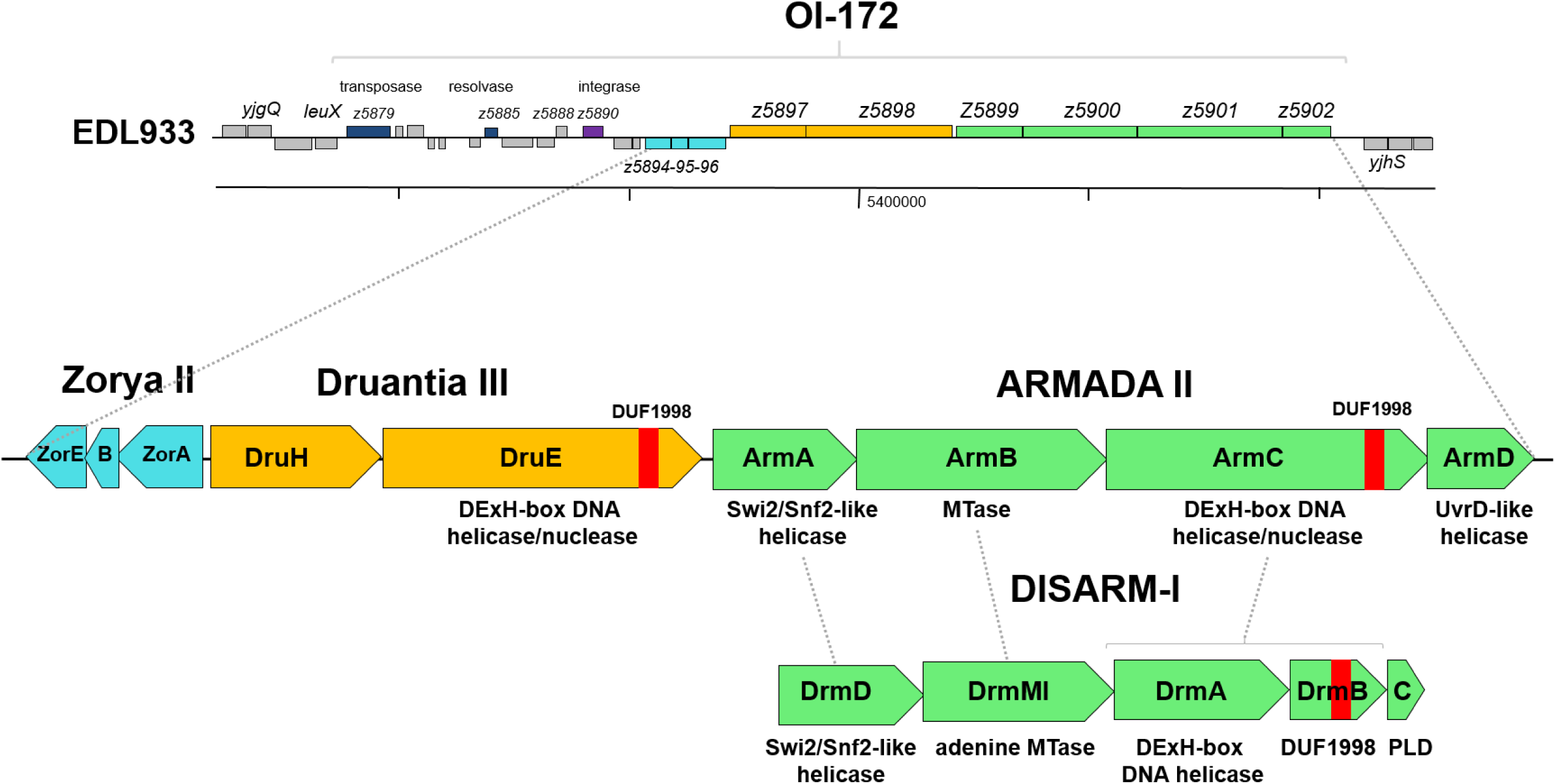
Gene organization of the phage defense island on OI-172 of EHEC O157:H7 strain EDL933 and the comparison to the DISARM I system. EDL933 encodes three phage defense systems on OI-172, i.e., Zorya II (*zorABE*), Druantia III (*druHE*), and the novel ARMADA II (*armABCD*) system, which is related to the DISARM I system of *Serratia* sp.. The homologies of ArmA, ArmB, and ArmC of ARMADA II (26) to the components of the DISARM I system (DrmMI, DrmD, DrmA, DrmB, and DrmC) are indicated by dashed lines. The two DExH box helicases/nucleases of the YprA family (26) (DruE and ArmC) contain a DUF1998 domain and resemble a DrmAB fusion with an additional DrmC-homologous PLD domain in DruE and a non-PLD domain in ArmC, respectively. ARMADA II additionally encompasses an UvrD-like NTPase/helicase (ArmD) that resembles the GajB subunit of the Gabija system (36).

The Zorya II-encoding *zorABE* operon (*z5894-z5896*) is divergently transcribed from the *druHE* operon and has been previously shown to protect against phages **(Fig. 1; Table S1)** (27, 28). Zorya II systems are distributed in EHEC O157:H7 and O145:H28 strains and in other pathogenic proteobacteria, including *Burkholderia* sp., *Vibrio* sp., *Serratia marcescens*, *Moraxella*, *Campylobacter*, *Legionella*, and *Pseudoalteromonas* sp. (25).

The third defense system of the OI-172 phage defense island, ARMADA II (26), is encoded by the *armABCD* operon (*z5899-z5902*) and transcribed downstream of the *druHE* operon **(Fig. 1; Table S1)**. Reminiscent of the Druantia III system, ARMADA II components show similarity to components of the DISARM I system (DrmD, DrmMI, DrmA, DrmB, DrmC) (22, 23, 25), with ArmA predicted as a DrmD-like Swi2/Snf2-type NTPase/helicase, ArmB as a DrmMI-like adenine MTase, ArmC as a DrmA-like DExH box helicase/nuclease fused to a DrmB-like DUF1998 domain protein, and two DrmC-like non-PLD nuclease domain proteins. ARMADA II additionally contains ArmD, a predicted UvrD-like NTPase/helicase resembling the GajB subunit of the Gabija defense system (36) **(Fig. 1; Table S1)**.

### Distribution of the OI-172 phage defense island across *E. coli* serotypes and pathotypes

KEGG homology analyses were used to investigate the distribution of the OI-172 phage defense island of *E. coli* EDL933 in bacteria. The genes encoding the defense systems Zorya II, Druantia III, and ARMADA II were found to co-localize particularly in atypical enteropathogenic *E. coli* (aEPEC) and STEC-aEPEC of serotypes O157:H7, O145:H28, and O55:H7, but also in the non-pathogenic antibiotic-resistant *E. coli* strains SMS-3-5 (37) and ATCC 8739 **(Table S1)**. Based on KEGG analyses, the three co-localizing defense systems were also found in *Enterobacter roggenkampii* 35734, *Pectobacterium polaris*, *Alteromonas macleodii* Balearic Sea AD45, and *Oleispira antarctica* **(Table S1)**.

Next, we aimed to analyze the prevalence of the OI-172 phage defense island across pathogenic and non-pathogenic *E. coli* serotypes, pathotypes, and phylogroups more systematically. Thus, we compared the genome sequence encoding the three defense systems of strain EDL933 against the NCBI RefSeq database, covering 3792 complete *E. coli* genome sequences of intestinal pathogenic and non-pathogenic strains (38). The genome results revealed the occurrence of the three co-localizing defense systems Zorya II, Druantia III, and ARMADA II in 426 *E. coli* strains, including non-pathogenic commensal *E. coli* and pathogenic aEPEC and STEC-aEPEC strains of various Clermont phylogroups comprising 66 distinct serotypes **(Table S2)**. Additionally, we identified 41 *E. coli* strains, which encode either Zorya II (n=17), Druantia III (n=1), ARMADA II (n=17), or combinations of two defense systems (n=6). The diversity of the complete defense island across the 426 *E. coli* genomes is mostly based on variations of the *armB* gene that encodes the adenine MTase of the ARMADA II system, whereas other components of the three defense systems are highly conserved (up-to 100% identity) as displayed in the phylogenetic tree **(Table S2; Fig. 2)**. Consistent with the KEGG analysis, the three defense systems were most prevalent in 277 STEC-aEPEC O157:H7 strains of phylogroup E, and also detected in STEC-aEPEC/ aEPEC O55:H7 (n=8) and O145:H28 (n=19) strains **(Table S2; Fig. 2)**. The three defense systems also co-localize in the genomes of 117 non-pathogenic *E. coli* of highly diverse phylogroups and serotypes, such as O15:H1, O17:H41, O45:H45, O83:H42, O146:H20, O153var1:H9/H34, and Onovel32:H9 as the most prevalent ones **(Table S2, Fig. 2)**. The bioinformatics analyses revealed that the defense islands show a lower degree of sequence conservation for the *armB* gene in non-pathogenic strains, forming diverse phylogenetic clusters distant from the O157/O55:H7 and O145:H28 lineages [EHEC1 clonal complex (39)] **(Table S2, Fig. 2)**. Overall, our data indicate that the defense island is distributed across many *E. coli* serotypes, phylogroups and lineages, but occurs most frequently and conserved in STEC-aEPEC of the EHEC1 clonal complex. Interestingly, other important human-pathogenic STEC/EHEC serotypes, such as O26:H11, O103:H2, O111:H8/H11, or O118:H16 [EHEC2 clonal complex (39)], completely lack the OI-172-associated defense island. Thus, the data suggest that the defense island has co-evolved with specific *E. coli* serotypes, such as the O157/O55 and O145 lineages, as well as diverse phylogroups and serotypes of non-pathogenic *E. coli*, but does not occur in the other STEC/EHEC lineage of the EHEC2 complex.

**Fig. 2.**
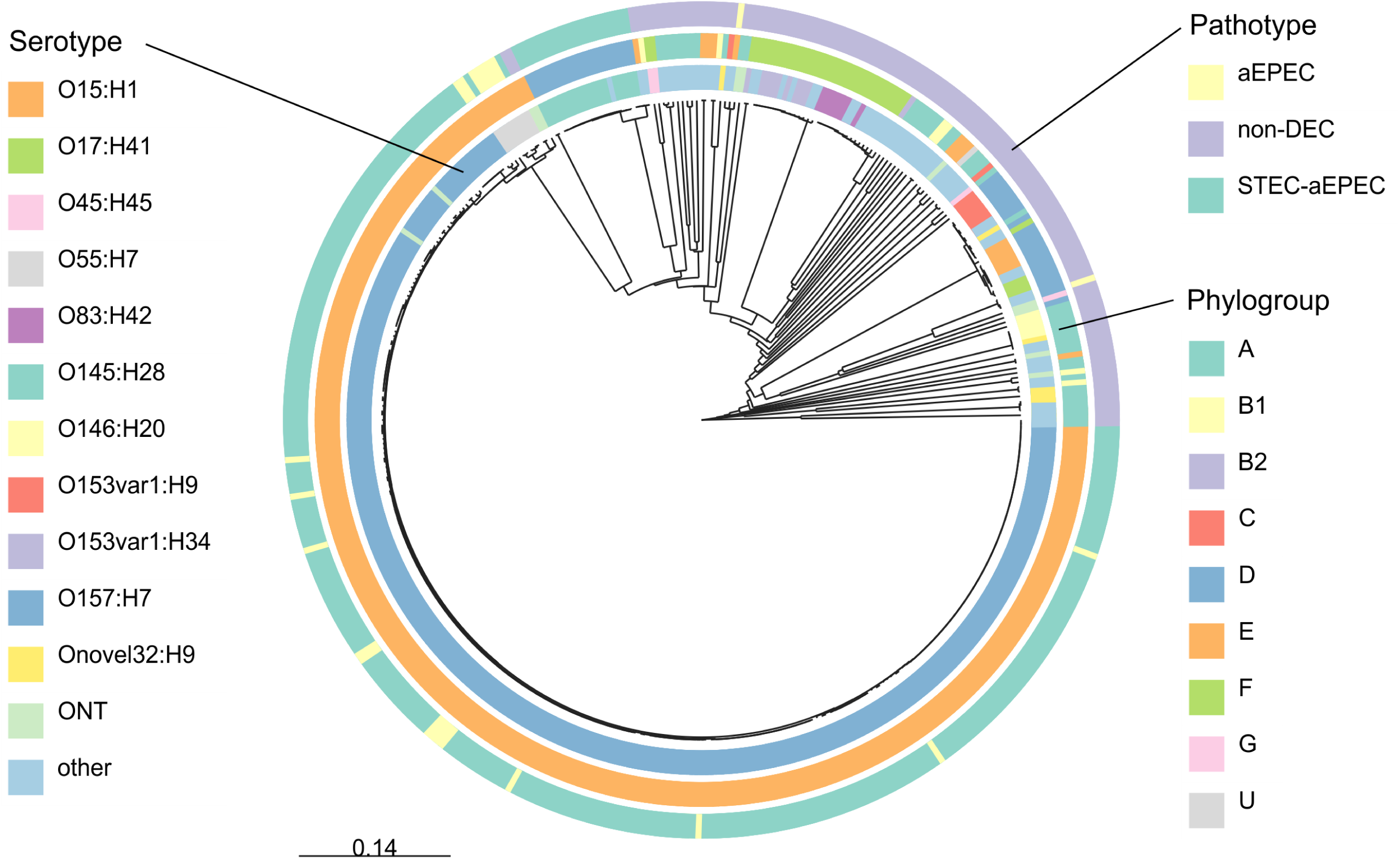
UPGMA tree of the three co-localizing phage defense systems Zorya II, Druantia III, and ARMADA II across 424 *E. coli* genomes, including the STEC-aEPEC serotypes O157:H7, O55:H7, and O145:H28 and diverse non-pathogenic strains. The tree was calculated based on the distance matrices of extracted genome regions of the co-localizing Zorya II, Druantia III, and ARMADA II defense system-encoding genes as retrieved from public *E. coli* genomes of the NCBI RefSeq database **(Table S2)** using mafft. Colors represent the respective strain characteristics serotype (inner circle, only serotypes with at least three strains were represented, ONT = not typeable), phylogroup (middle circle), and pathotype (outer circle, aEPEC = atypical enteropathogenic *E. coli*, STEC = Shiga toxin-producing *E. coli*, noDEC = no diarrheagenic *E. coli*).

Overall, the co-localization of the three defense systems in several STEC-aEPEC strains points to their major roles in the defense against a broad spectrum of phages that attack these enteropathogenic bacteria. Moreover, the homologies between the Druantia III and ARMADA II-associated DExH box helicases/nucleases DruE and ArmC suggest additive and synergistic activities in the phage defense to provide more robust immunity of EHEC strains against various phages than the individual systems alone.

### The EHEC strain EDL933 Δ*stx1/2* is resistant to many phages of the BASEL phage collection

We hypothesized that the three OI-172-encoded phage defense systems confer a broad range of phage resistance in the EHEC strain EDL933. Thus, we selected a panel of 44 *E. coli* phages of the BASEL phage collection (34), belonging to different phage families, including *Drexlerviridae, Siphoviridae, Demerecviridae, Tevenvirinae, Vequintavirinae, Autographiviridae* and temperate phages, to determine the phage susceptibility of the EHEC strain EDL933 Δ*stx1/2* by using plaque-forming unit (PFU) assays. For comparison of phage susceptibility of the EHEC strain, we used the *E. coli* K-12 laboratory strain W3110 in PFU assays **(Fig. 3; Table S3)**. Based on the PFU assays, we calculated the EOP as log_10_-fold changes of average PFU values of the EHEC strain versus *E. coli* W3110 to determine the phage susceptibility of EHEC. The results showed that the W3110 strain is very susceptible to the 44 phages, whereas the EHEC strain is highly resistant, as revealed by EOP differences of 10^6^-fold for the EHEC strain versus the W3110 strain **(Fig. 3; Table S3)**. Specifically, the EHEC strain exhibited complete resistance against *Drexlerviridae, Siphoviridae, Tevenvirinae, Vequintavirinae* and *Autographividae*. Only phages of the *Demerecviridae* were able to efficiently lyse the EHEC strain, including *Epseptimaviruses* (Bas26, Bas27, Bas28, Bas29, Bas30) and *Tequintaviruses* (Bas32, Bas33, Bas34). Here we observed EOP differences of 10^2^-10^3^-fold between the EHEC strain and *E. coli* W3110, indicating that *Demerecviridae* are successful anti-EHEC phages. Altogether, our results clearly point to strong resistance of the EHEC strain EDL933 Δ*stx1/2* towards a broad range of different phage families.

**Fig. 3.**
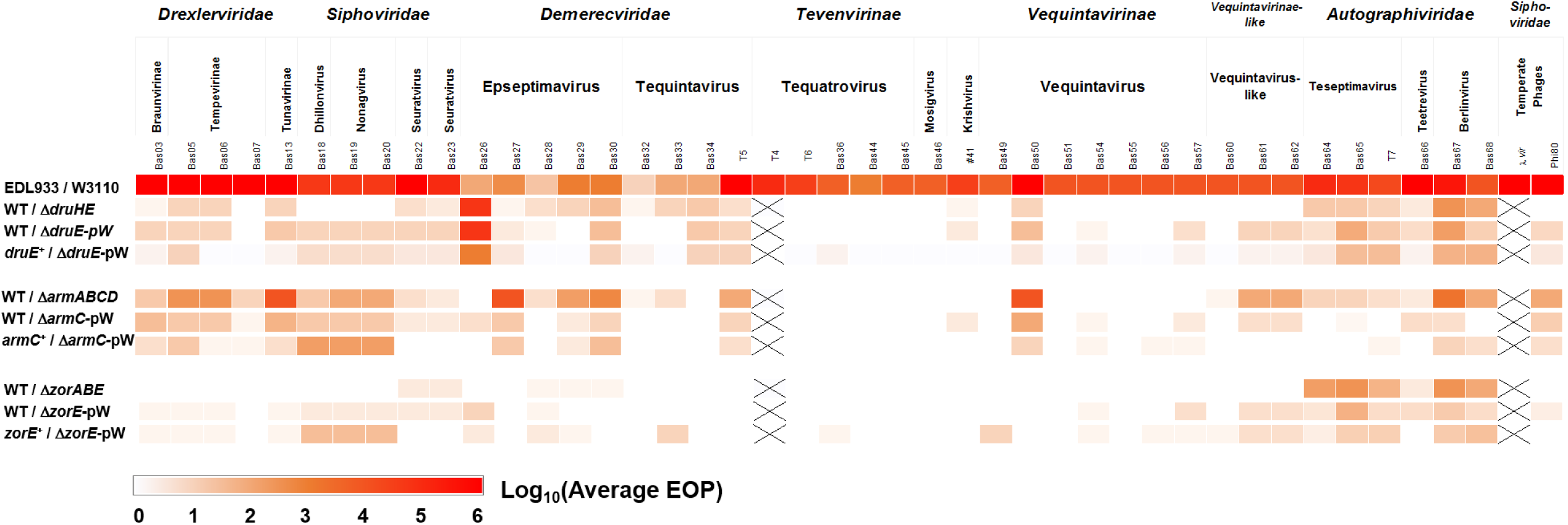
The OI-172-encoded Zorya II, Druantia III, and ARMADA II defense systems confer resistance against a broad phage spectrum in strain EDL933 Δ*stx1/2*. Heatmap showing protection by the defense systems against a spectrum of 44 phages. First, the phage resistance of EHEC versus the sensitive *E. coli* W3110 was determined in PFU/EOP assays. Second, the log_10_ fold-changes in phage defense systems were determined using PFU/EOP assays of the 44 phages in the EDL933 Δ*stx1/2* (WT) strain versus the Δ*zorABE*, Δ*druHE*, or Δ*armABCD* deletion mutants. Third, PFU/EOP assays were used to validate the complementation of the Δ*zorE,* Δ*druE*, and Δ*armC* effector mutants with pWKS30-encoded z*orE, druE*, *and armC* (z*orE^+^, druE^+^*, *and armC^+^*) versus the mutants with empty pWKS30 plasmids (Δ*zorE-*pW, Δ*druE*-pW, Δ*armC*-pW). Log_10_ EOP values are averages from 3-4 biological replicate experiments, which are presented in **Tables S3-S5**. The color code below the heatmap indicates the scale of log_10_ EOP values.

### The Zorya II, Druantia III, and ARMADA II phage defense systems on OI-172 provide a broad range of anti-phage defense against overlapping and specific phages

To analyze the phage spectrum targeted by the three phage defense systems, we constructed individual Δ*zorABE*, Δ*druHE*, or Δ*armABCD* deletion mutants lacking the Zorya II, Druantia III, or ARMADA II systems in *E. coli* EDL933 Δ*stx1/2*. EOP assays were used to analyze the individual defense system mutants for their phage susceptibility against our panel of 44 BASEL phages (34) in comparison to *E. coli* EDL933 Δ*stx1/2*, which is referred to as wild type (WT) **(Fig. 3; Table S4)**. Additionally, we constructed Δ*zorE,* Δ*druE*, and Δ*armC* single mutants deficient for each helicase/nuclease effector, as well as the corresponding complemented strains, expressing *zorE, druE*, and *armC* ectopically from plasmid pWKS30 under the IPTG-inducible *lac* promoter in the respective defense mutants in the background of strain EDL933 Δ*stx1/2*.

The EOP assays of the Δ*zorABE*, Δ*druHE*, and Δ*armABCD* mutants revealed that the three defense systems contribute to different extents to the defense against specific and overlapping phages and different phage families. Specifically, the *zorABE* mutant exhibited 10^2^-10^3^-fold enhanced sensitivity towards the *Autographiviridae*, but very weak 10-fold differences in phage susceptibilities to *Seuratviruses* (Bas22, Bas23) **(Fig. 3; Table S4)**. The EOP assays of the *zorE* mutant versus the WT largely confirmed this spectrum of phage protection, with very weak differences in phage susceptibility against *Siphoviridae* and stronger protection of Zorya II against *Autographiviridae*. Moreover, complementation studies with pWKS30-*zorE* restored the resistance of the Δ*zorE* mutant against podoviruses and *Siphoviridae* (Bas18-20) to WT level **(Fig. 3; Table S5)**. These results indicate that Zorya II protects mainly against the small podoviruses of *Autographiviridae*.

Deletion of Druantia III in strain EDL933 Δ*stx1/2* led only to weakly, 10-10^2^-fold enhanced phage susceptibility towards several phages of *Drexlerviridae* (Bas5, 6)*, Siphoviridae* (Bas22, Bas23)*, Demerecviridae* (Bas28, Bas29, Bas30, Bas33, Bas34, T5) and *Vequintavirinae* (Bas50) **(Fig. 3; Table S4)**. However, the Δ*druHE* mutant showed higher, 10^2^-10^4^-fold phage sensitivity against the *Autographiviridae* and the *Epseptimavirus* Bas26, as compared to *E. coli* EDL933 Δ*stx1/2* (WT). In contrast, the *druHE* mutant retained its resistance against *Tevenvirinae*, including *Tequatroviruses* and *Mosigvirus*. The EOP assays using the Δ*druE* effector mutant versus the WT confirmed the pattern of phage protection as observed for the Δ*druHE* mutant, showing increased susceptibilities against phages of *Drexlerviridae, Siphoviridae, Demerecviridae, Vequintavirinae*, and *Autographiviridae* **(Fig. 3; Table S5)**. Complementation with pWKS30-*druE* successfully restored the phage resistance of the Δ*druE* mutant to WT levels **(Fig. 3; Table S5)**. These results show that Druantia III provides weak immunity against a broad spectrum of phages of various phage families, with stronger protection against selected phages, such as Bas26 and *Autographiviridae*. However, Zorya II and Druantia III do not protect against *Tevenvirinae* (*Tequatroviruses* and *Mosigviruses*) due to their hypermodified phage genomes, confirming previous findings (35). Interestingly, the EOP assays of the Δ*armABCD* mutant, deficient in ARMADA II, showed strongly enhanced sensitivities against a broad spectrum of phages belonging to the *Drexlerviridae, Siphoviridae, Demerecviridae, Vequintavirinae*, and *Autographiviridae* **(Fig. 3; Table S4)**. Moreover, the phage sensitivity of the Δ*armABCD* mutant was much higher compared to the Δ*druHE* mutant, indicating that ARMADA II and Druantia III might act additively or synergistically against a similar phage spectrum. Specifically, the Δ*armABCD* mutant showed 10^2^-10^5^-fold higher phage sensitivity versus the WT against *Drexlerviridae* and *Siphoviridae*, whereas Druantia III conferred only very weak protection against a few phages of these phage families **(Fig. 3; Table S4)**. Similarly, a high susceptibility of the Δ*armABCD* mutant of 10^3^-10^5^-fold versus the WT was observed against some *Epseptimaviruses* (Bas27, Bas29, Bas30), *Tequintavirus* (T5), and *Vequintaviruses* (Bas50, Bas61, Bas62), while the Δ*druHE* mutant showed only slightly increased phage sensitivity (10^0^-10^1^-fold) **(Fig. 3; Table S4)**. However, the EOP assays revealed a similar 10^2^-10^4^-fold higher phage sensitivity of the Δ*armABCD* and Δ*druHE* mutants versus the WT against the *Autographiviridae* **(Fig. 3; Table S4)**. Furthermore, EOP assays with the Δ*armC* mutant, lacking the DExH-box DNA helicase/nuclease effector, resulted in enhanced sensitivity against phages of the *Drexlerviridae*, *Siphoviridae*, *Vequintavirinae*, and *Autographiviridae* **(Fig. 3; Table S5)**. Complementation of the Δ*armC* mutant with pWKS30-*armC* restored resistance against several phages, albeit at a lower level than observed for the WT, indicating incomplete complementation with ArmC using the IPTG-inducible pWKS30 plasmid in the EHEC strain **(Fig. 3; Table S5)**. Overall, our results indicate that ARMADA II provides the strongest level of immunity against a broad spectrum of phages, as compared to Zorya II and Druantia III, while each of these defense systems similarly contributes to resistance towards *Autographiviridae*. However, none of these nuclease-based defense systems confers protection against the *Tevenvirinae*, including *Tequatroviruses* and *Mosigvirus* **(Fig. 3; Table S4)**, which are known to have hypermodified phage genomes (40, 41). Thus, consistent with their resistance against DNA-targeting RM systems (34), *Tevenvirinae* also showed complete resistance against Zorya II, Druantia III, and ARMADA II of this EHEC-associated phage defense island.

### The individual components of the Zorya II, Druantia III, and ARMADA II systems are essential for the full anti-phage defense

After having elucidated the susceptible phage spectrum for the three phage defense systems, we investigated whether the individual components of the Zorya II, Druantia III, and ARMADA II systems are required for the specific anti-phage defense. Thus, we used growth assays in liquid medium to study the population phenotype of the single and multiple defense component mutants for each of the three defense systems, including Δ*zorA,* Δ*zorB,* Δ*zorE*, and Δ*zorABE* mutants for the Zorya II system, Δ*druH,* Δ*druE,* and Δ*druHE* mutants for the Druantia III system and Δ*armA,* Δ*armB,* Δ*armC,* Δ*armD* and Δ*armABCD* mutants for the ARMADA II system after infection with a susceptible phage at a specific multiplicity of infection (MOI) **(Fig. 4A-C)**. Based on our EOP assays, we selected T7 as a susceptible phage for Zorya II, Bas26 for Druantia III, and Bas18 for ARMADA II in liquid growth assays. The EDL933 Δ*stx1/2* defense mutants, deficient in single and multiple components of the defense systems, were grown in Luria broth (LB) and infected with the susceptible phages at an OD_600_ of 0.08 to monitor the growth and collapse of the cultures upon phage propagation.

**Fig. 4.**
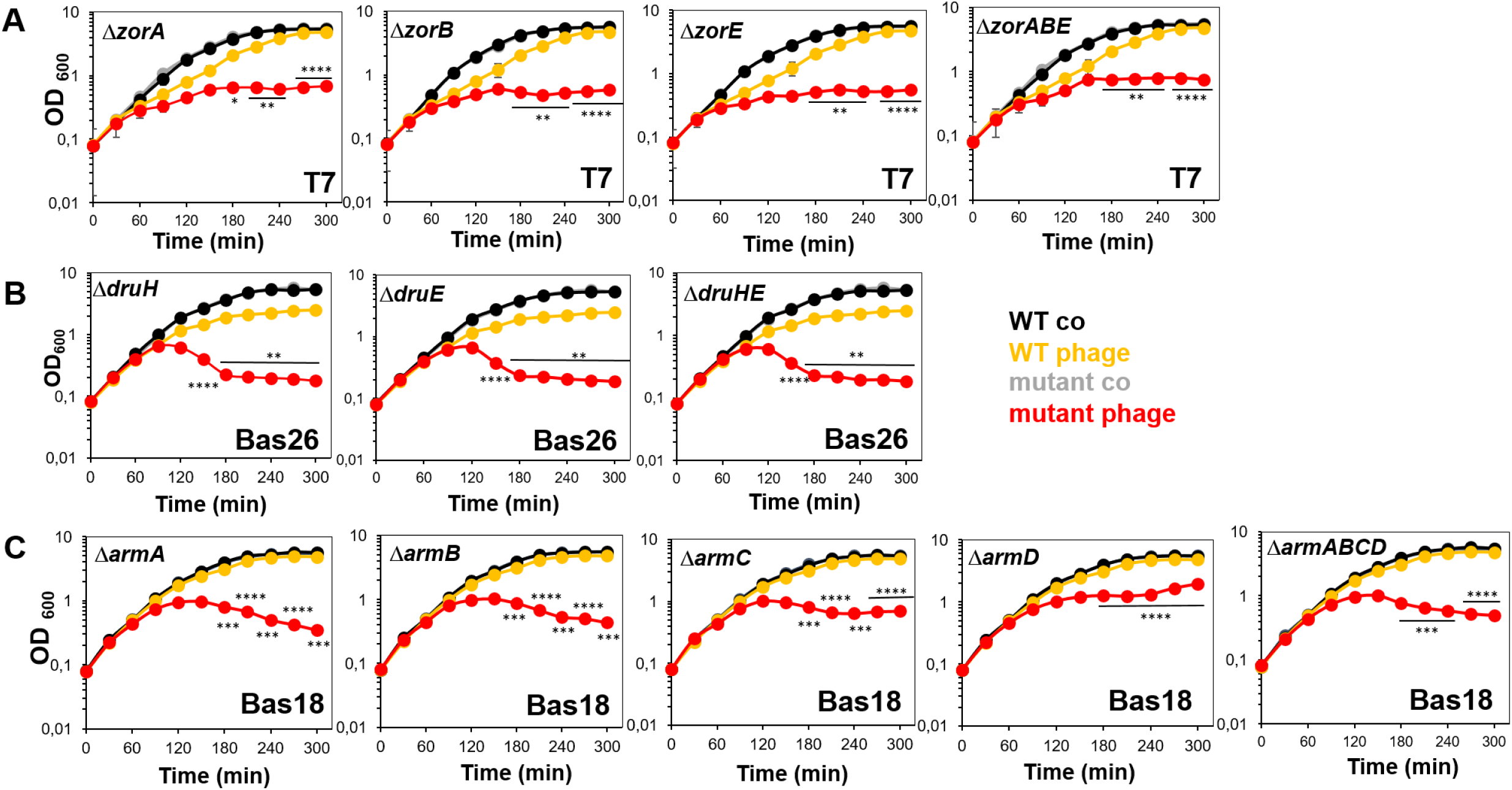
The individual components of the Zorya II, Druantia III, and ARMADA II systems are essential for the defense against sensitive phages (T7, Bas26, Bas18). Growth curves of the strain EDL933 Δ*stx1/2* (WT) (yellow) versus **(A)** the Δ*zorA,* Δ*zorB,* Δ*zorE*, and Δ*zorABE* mutants (red) after infection with phage T7 (MOI 7), **(B)** the Δ*druH,* Δ*druE*, and Δ*druHE* mutants (red) after phage Bas26 infection (MOI 3), and **(C)** the Δ*armA*, Δ*armB*, Δ*armC*, Δ*armD*, and Δ*armABCD* deletion mutants (red) after infection with phage Bas18 (MOI 7). The EHEC strains were grown in LB until an OD_600_ of 0.08 and subjected to phage infection to monitor the growth and collapse of the bacterial culture. The growth of the strains without phage infection was also monitored, revealing no differences between WT (black) and defense mutants (grey). The experiments were performed in 3-4 biological replicates. The data points indicate average values, and error bars represent the standard deviations (SD). *P* values of infected WT vs. mutants indicate: *, *p* < 0.05; **, *p* < 0.01; ***, *p* < 0.001.

The growth assays of the WT, Δ*zorA,* Δ*zorB,* Δ*zorE*, and Δ*zorABE* mutants after infection with phage T7 (MOI 7) showed that all Zorya mutants were sensitive and unable to grow upon phage infection, while the growth of WT was only slightly affected **(Fig. 4A)**. These data confirmed previous findings that the ZorA, ZorB, and ZorE components are essential for the Zorya II-mediated anti-phage defense (25).

After infection of the WT, the Δ*druH,* Δ*druE*, and Δ*druHE* mutants with Bas26 (MOI 3), all strains continued with growth until an OD_600_ of 0.6, followed by a collapse of the mutant cultures, whereas the WT was resistant and able to grow at a slightly reduced growth rate **(Fig. 4B)**. These results indicate that both proteins DruH and DruE are essential for Druantia III-mediated phage resistance.

Similarly, the strain EDL933 Δ*stx1/2* (WT) was resistant to infection with Bas18 (MOI 7) and continued to grow as the non-treated control. However, Bas18-infected Δ*armA,* Δ*armB,* Δ*armC,* Δ*armD*, and Δ*armABCD* mutants only grew to an OD_600_ of 0.08, followed by lysis of the strains **(Fig. 4C)**. Of note, the growth of the Δ*armD* mutant was less affected after Bas18 infection compared to the Δ*armA,* Δ*armB*, or Δ*armC* mutants, suggesting a reduced requirement of the NTPase/helicase ArmD for the ARMADA II-dependent anti-phage defense.

### Synergy between the Zorya II, Druantia III, and ARMADA II systems in the anti-phage defense of the EHEC strain EDL933 Δ*stx1/2*

The modularity of the co-occurring Zorya II, Druantia III, and ARMADA II defense systems suggests their synergistic interactions and complementary activities in the anti-phage defense in the EHEC host strain EDL933 Δ*stx1/2*. Thus, we deleted the co-localizing defense system pairs Zorya II/ Druantia III and Druantia III/ ARMADA II as well as the three systems Zorya II/ Druantia III/ ARMADA II in strain EDL933 Δ*stx1/2*. Using EOP assays, the phage sensitivities of the Δ*zor-dru*, Δ*dru-arm*, and Δ*zor-dru-arm* combined system mutants were analyzed in comparison to the single system mutants against the panel of 44 phages **(Fig. 5; Table S4)**. To assess synergistic interactions, the synergy score was defined as EOP values greater than the sum of the combined anti-phage effects of the single defenses, which is displayed as a yellow-green color code in the heatmap **(Fig. 5; Table S4)**. Specifically, yellow and green synergy scores indicate that the EOP of the combined system mutant (e.g., Δ*zor-dru* or Δ*dru-arm*) exceeds values of 10^1^ and 10^2^ over the additive EOP of the single system mutants, respectively **(Fig. 5; Table S4)**.

**Fig. 5.**
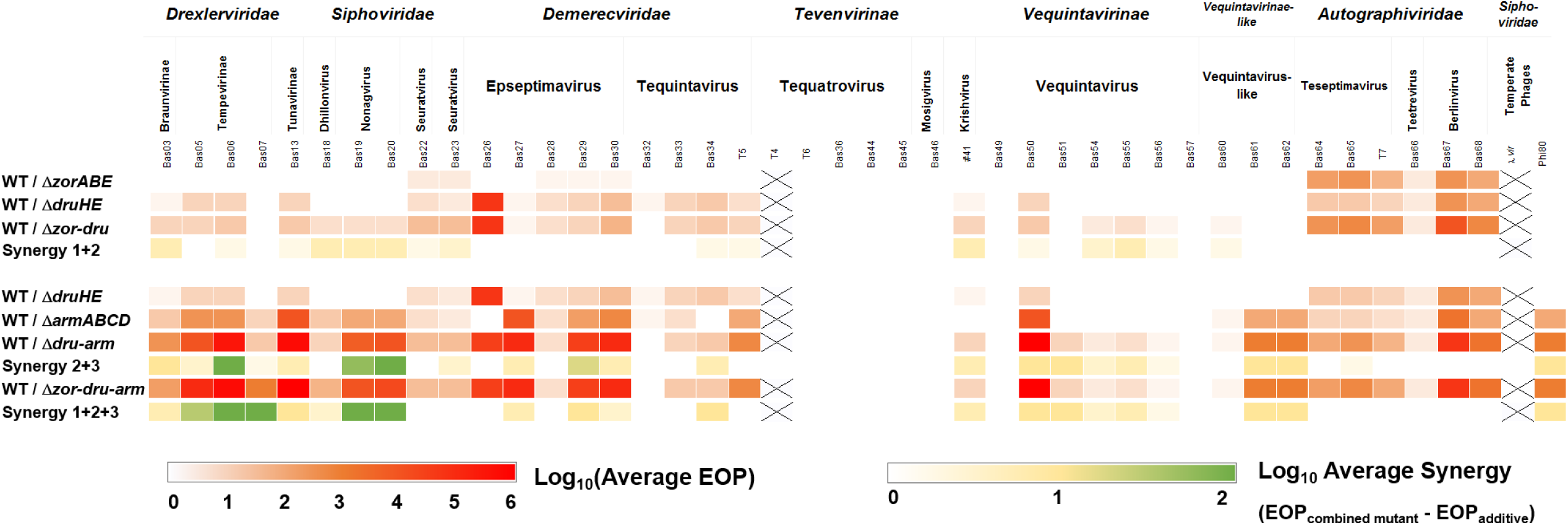
The Druantia III and ARMADA II defense pair provides strong synergistic anti-phage activity. The heatmap shows the log_10_ EOP values of 44 phages in the Δ*zorABE,* Δ*druHE*, and Δ*armABCD* single system mutants and the combined Δ*zor-dru*, Δ*dru-arm*, and Δ*zor-dru-arm* mutants quantified versus *E. coli* EDL933 Δ*stx1/2* (WT). The yellow-green synergy score indicates the log_10_ higher EOP values of the combined Δ*zor-dru,* Δ*dru-arm*, or Δ*zor-dru-arm* mutants above the calculated additive protection by two or three single-system mutants. Log_10_ EOP values are averages from 3-4 biological replicate experiments, which are presented in **Tables S3-S5**.

Overall, the results of the EOP assays showed that Zorya II and Druantia III act only very weakly synergistically against *Drexlerviridae* (Bas03, Bas06), *Siphoviridae* (Bas18, Bas19, Bas20, Bas23), *Krishvirus* and *Vequintavirinae* (Bas50, Bas54, Bas55), since the EOP of the Δ*zor-dru* mutant exceeded values of <10^1^ over the additive effects of the single system mutants **(Fig. 5; Table S4)**. As Zorya II alone was not effective against the *Epseptimaviruses* Bas28, Bas29, and Bas30, no synergy of Zorya II and Druantia III was observed against these phages in the EOP assay. Interestingly, while Δ*zorABE* and Δ*druHE* single system mutants showed strongly enhanced sensitivity towards *Autographiviridae*, the corresponding sensitivity of the combined Δ*zor-dru* mutant only slightly exceeded that of the Δ*zorABE* single mutant **(Fig. 5; Table S4)**. These results indicate that Zorya II and Druantia III act weakly synergistically against those phages, for which the single system provided weak protection, whereas no additive or synergistic defense was observed against *Autographiviridae*, which are strong targets of both individual systems.

We further used EOP assays to analyze the sensitivity of the combined Δ*dru-arm* mutant versus the Δ*druHE* and Δ*armABCD* single system mutants to reveal possible synergistic interactions between Druantia III and ARMADA II. The combined deletion of Druantia III and ARMADA II resulted in hypersensitivity towards many phages of the *Drexlerviridae, Siphoviridae, Demerecviridae*, and *Vequintavirinae*, as compared to the single system mutants **(Fig. 5; Table S4)**. Specifically, the EOP of the Δ*dru-arm* mutant exceeded values of up to 10^2^ over the sum of the EOP values determined for the single system mutants. These data point to a synergistic interaction of Druantia III and ARMADA II in the defense against several phages, such as Bas6, Bas19, and Bas20 with the highest synergy score of 10^2^ over the additive EOP. Additionally, Druantia III and ARMADA II provide additive protection against *Autographiviridae*, since the Δ*dru-arm* mutant displayed enhanced sensitivity, but no hypersensitivity, compared to the single mutants. However, in the Δ*zor-dru-arm* full defense mutant, we only observed a minor increase in the phage susceptibility against a few phages (e.g., Bas5, Bas7, and Bas18), in comparison to the Δ*dru-arm* mutant **(Fig. 5; Table S4)**. Taken together, these data strongly support that Druantia III and ARMADA II act additively and synergistically in the defense against a broad spectrum of phages, together providing more robust innate immunity than the individual systems. Moreover, the presence of Druantia III and ARMADA II is sufficient to confer broad specificity and strong anti-phage defense, while Zorya II additionally protects against *Autographiviridae*.

To validate our results from the EOP assays, we used growth assays of the EDL933 Δ*stx1/2* combined system mutants (Δ*zor-dru*, Δ*dru-arm* and Δ*zor-dru-arm*) in comparison to the single system mutants. The defense mutants were infected at OD_600_ 0.08 with phages against which Zorya II, Druantia III and ARMADA II provide strong and synergistic defense, as revealed by high EOP values of single system mutants and a high synergy score of combined mutants **(Fig. 6; Fig. S1)**. Specifically, we selected Bas6 (*Tempervirinae*), Bas20 (*Nonagvirus*), Bas26, Bas27, Bas30 (*Epseptimaviruses*), Bas50 (*Vequintavirus*), and Bas67 (*Berlinvirus*) as most susceptible phages for the three defense systems to monitor the growth of the EDL933 Δ*stx1/2* defense mutants after phage infection at different MOIs **(Fig. 6; Fig. S1)**. For quantification of the synergy from the growth curves after phage infection, we calculated the area under the curve (AUC) for different MOIs of the single and combined system mutants **(Fig. 6; Fig. S1)**. The AUC values of the defense mutants were normalized to the uninfected control, which was set to 100%. Synergistic effects of the defense pairs Zorya II/Druantia III and Druantia III/ARMADA II were considered when the %AUC values of the combined system mutants were <0.5-fold lower compared to the most sensitive single system mutant.

**Fig. 6.**
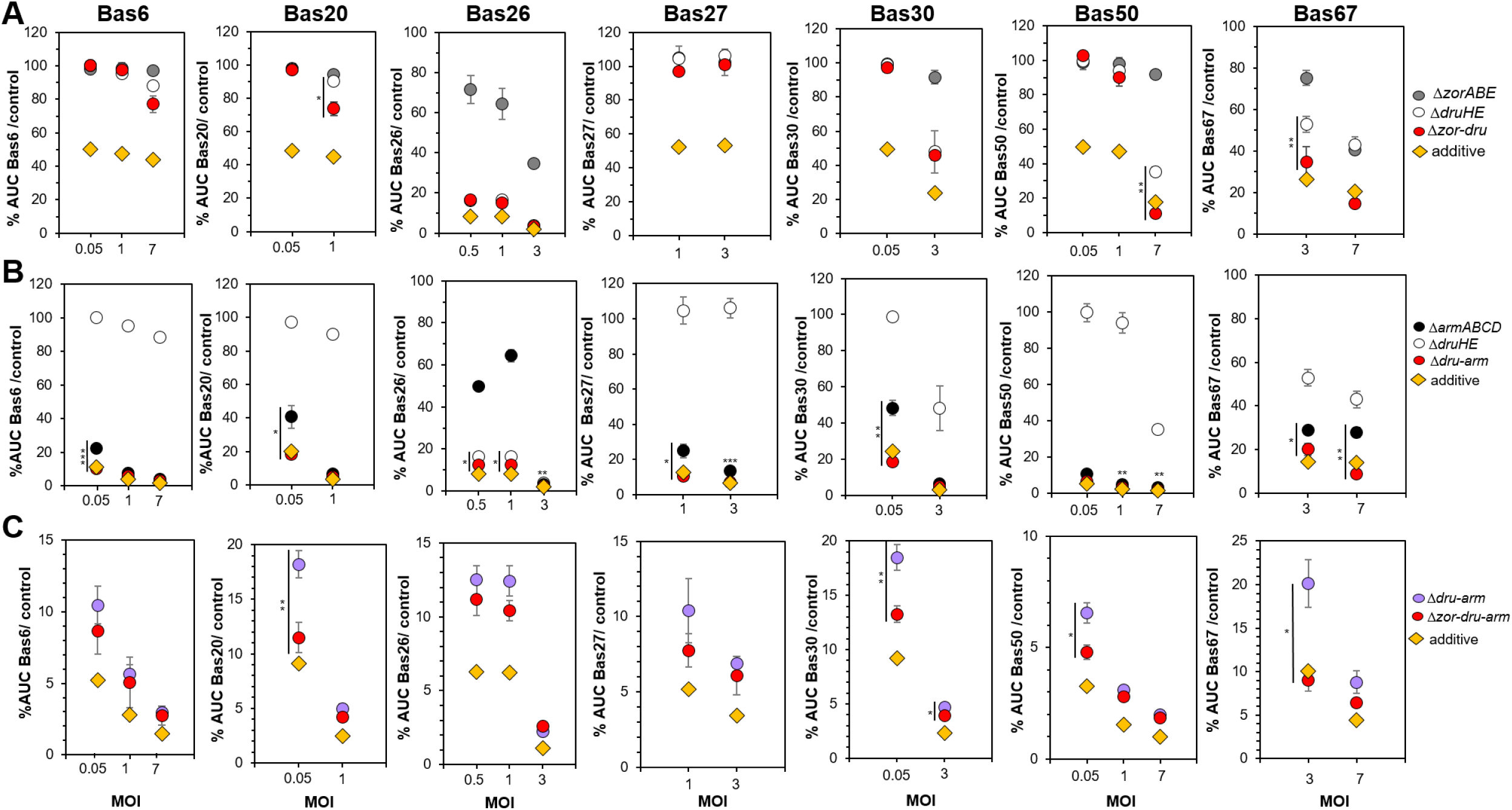
AUC values of growth curves support additive interactions of Druantia III and ARMADA II at the population level. Growth curves were recorded for the Δ*zorABE,* Δ*druHE*, and Δ*armABCD* single system mutants and the combined Δ*zor-dru*, Δ*dru-arm*, and Δ*zor-dru-arm* mutants in LB after infection with different phages (Bas6, Bas20, Bas26, Bas27, Bas30, Bas 50, and Bas 67) at different MOIs **(Fig. S1A-G)**. Average values of three biological replicates of the growth curves for the defense mutants are presented for the seven phages with error bars indicating the SD **(Fig. S1A-G)**. The AUC was calculated from the phage-infected culture and normalized to that of the uninfected control, which was set to 100%. The combined defenses of Zorya II and Druantia III **(A)**, Druantia III and ARMADA II **(B)**, or of all three systems **(C)** are synergistic when the %AUC value of the combined mutant (red dot) is below the additive value (yellow square). The additive value was defined as 0.5-fold of the %AUC value of the most sensitive single system mutant **(A, B)** or of the Δ*dru-arm* double mutant **(C)**. *P* values indicate significant %AUC differences of the infected combined system mutant versus the most sensitive single mutant **(A, B)** or versus the *dru-arm* mutant **(C)**: *, *p* < 0.05; **, *p* < 0.01; ***, *p* < 0.001.

The growth and AUC comparisons revealed that the Δ*zorABE,* Δ*druHE*, and Δ*zor-dru* mutants were similarly resistant to the phages Bas6, Bas20, Bas27, and Bas30 (MOI 0.05-3) as reflected by high %AUC values, indicating no additive or synergistic effects of the combined Zorya II/ Druantia III systems **(Fig. 6; Fig. S1)**. Thus, the weak synergy of Zorya II/ Druantia III observed against Bas6 and Bas20 in EOP assays could not be validated at the population level. In EOP assays, Bas26 was identified as the strongest target of Druantia III **(Fig. 3; Table S4)**. In growth assays, the Δ*druHE* and Δ*zor-dru* mutants were equally sensitive to Bas26 (MOI 0.05) with collapse of the culture and low %AUC values with no differences, revealing no additive effects of the Zorya II/Druantia III defenses **(Fig. 6; Fig. S1)**. However, the Zorya II/ Druantia III defense pair conferred synergistic protection against Bas50 and Bas67 (MOI 7), since the %AUCs of the Δ*zor-dru* mutant were <0.5-fold decreased compared to the most susceptible Δ*druHE* single system mutant **(Fig. 6; Fig. S1)**. While no synergistic effects of Zorya II/ Druantia III against *Autographiviridae* were detected in the EOP assays, the growth results provide additional evidence that the Zorya II/ Druantia III systems act also synergistically against podoviruses, such as Bas67, at the population level **(Fig. 6; Fig. S1)**.

The EOP assays had revealed very strong synergy between Druantia III and ARMADA II against a broad spectrum of different phage families **(Fig. 5; Table S4)**. To validate the interactions between the Druantia III/ ARMADA II defenses at the population level, we analyzed the growth of the combined Δ*dru-arm* mutant versus the Δ*druHE* and Δ*armABCD* single system mutants **(Fig. 6; Fig. S1)**. Consistent with the EOP assays, the Δ*armABCD* mutant was much more sensitive in growth after infection with Bas6, Bas20, Bas27, Bas30, Bas50, and Bas67 than the Δ*druHE* mutant, and this growth defect was further pronounced in the combined Δ*dru-arm* mutant. Specifically, the %AUC values of the Δ*dru-arm* mutant were ∼0.5-fold reduced after infection with Bas6, Bas20, Bas50 (MOI 0.05) and Bas27 (MOI 1), compared to the Δ*armABCD* single system mutant, indicating additive interactions of the Druantia III/ ARMADA II defenses **(Fig. 6; Fig. S1)**. Infection of the Δ*dru-arm* mutant with Bas30 (MOI 0.05) and Bas67 (MOI 7) resulted in lower %AUC values than the additive effects, pointing to synergistic interactions of the co-occurring defense systems **(Fig. 6; Fig. S1)**. However, similarly strong sensitivities in growth with little differences in %AUC values were observed for the Δ*dru-arm* and Δ*druHE* mutants upon infection with Bas26, indicating that protection comes exclusively from Druantia III **(Fig. 6; Fig. S1)**. Taken together, the growth data revealed synergistic effects of the Druantia III/ARMADA II combined defenses against Bas30 and Bas67 as well as additive effects against Bas6, Bas20, Bas27, and Bas50 **(Fig. 6; Fig. S1)**. In contrast, EOP assays showed strongly synergistic effects of Druantia III and ARMADA II against Bas6 and Bas20. Moreover, both defense pairs, Zorya II/ Druantia III and Druantia III/ ARMADA II, conferred synergistic protection against Bas67 of the *Autographiviridae* at the population level, whereas only additive effects could be quantified for the Druantia III/ ARMADA II pair in the EOP assays.

The growth comparison of the Δ*zor-dru-arm* versus Δ*dru-arm* mutants revealed that the deletion of all three defense systems, Zorya II, Druantia III, and ARMADA II, led to a further decrease of the %AUC values after phage infection, with significant changes for Bas20, Bas30, Bas50, and Bas67 at low MOIs and non-significant changes for Bas6, Bas26, and Bas27 **(Fig. 6; Fig. S1)**. However, only for Bas67, the changes in %AUC values between Δ*zor-dru-arm* and Δ*dru-arm* mutants were in the range of additive protection, indicating that Zorya II additively contributes towards phage defense against the *Autographiviridae*. Overall, our EOP and growth results revealed additive and synergistic interactions between the Druantia III/ ARMADA II defenses against a broad phage panel, whereas Zorya II contributes with additive and synergistic effects against *Autographiviridae* in the EHEC strain EDL933 Δ*stx1/2*.

## DISCUSSION

In this manuscript, we discovered an OI-172-associated phage defense island of EHEC strain EDL933 Δ*stx1/2*, which encodes the three nuclease-based defense systems, Zorya II, Druantia III, and ARMADA II. This phage defense island is most prevalent in the STEC-aEPEC serotypes O157:H7, O145:H28, and O55:H7 (EHEC1 clonal complex), but also occurs in diverse serotypes and phylogroups of non-pathogenic *E. coli* **(Fig. 2; Table S2)**. The STEC O157:H7 lineage has evolved from an O55:H7 ancestor (42–44). Additionally, STEC O145:H28 strains are evolutionarily related to the O157:H7/O55:H7 lineage, since O145:H28 diverged as a sub-lineage before the separation of O157 from O55 (45). Thus, it appears that the evolution of the phage defense island is related to the O55:H7/O157:H7/O145:H28 lineage, whereas other more prevalent STEC EHEC serogroups, such as O26, O103, O111, or O118 of the EHEC2 clonal complex, do not harbor the phage defense island (45) **(Table S2)**. It is interesting to note that the STEC-aEPEC O157/O55/O145 serotypes carry up to 18 prophages and prophage-like elements (42–45). Thus, this phage defense island might have evolved together with their encountered phage foes, which were integrated into the genomes as lambdoid prophages.

Predictions using DefenseFinder revealed that *E. coli* genomes encode, on average, 5 to 7 defense systems (20, 29–31). Apart from the OI-172-encoded defense systems, *E. coli* O157:H7 strains of phylogroup E2 encode the abundant restriction modification systems RM I, RM II, RM IIG, and CRISPR-Cas type I-E among others (31). Previous studies revealed a correlation of the number of defense systems with phage resistance phenotypes in clinical and antibiotic-resistant *P. aeruginosa* isolates (32, 33). Here, we report that the EHEC outbreak strain EDL933 Δ*stx1/2* is highly resistant towards the majority of phage families of the BASEL collection, including *Drexlerviridae, Siphoviridae*, *Demerecviridae, Myoveridae (Tevenvirinae, Vequintavirinae)*, and *Autographiviridae* with EOP differences of 10^6^-fold compared to the phage-susceptible *E. coli* K-12 strain W3110. However, the EHEC strain was less resistant towards *Epseptimaviruses* and *Tequintaviruses*, as revealed by lower EOP differences of 10^2^-10^3^ compared to the *E. coli* strain W3110. Consistent with our results, O157:H7 strains were efficiently lysed by other *Epseptimaviruses*, which bind to the BtuB outer membrane protein as a secondary host cell receptor (14–16, 34).

In this study, we further investigated the contribution of the co-localizing Zorya II, Druantia III, and ARMADA II defense systems on OI-172 towards phage resistance of EHEC strain EDL933 Δ*stx1/2*. Our EOP results showed that Zorya II confers strong protection against *Autographiviridae* (*Teseptimavirus, Teetrevirus*, and *Berlinvirus*), but not against other phage families. These results confirm previous findings of plasmid-expressed Zorya II in *E. coli* strains MG1655 and MT56, providing strong immunity against phage T7, as indicated by EOP differences of 10^3^ compared to the *E. coli* strain with an empty plasmid (25, 27, 28). However, in contrast to previous work with the phage-sensitive strains MG1655 and MT56, the EDL933 Δ*stx1/2* WT strain was highly resistant against most phages, including T7 phages. Thus, at the population level, we only observed sensitive growth phenotypes of the Δ*zorABE* mutant upon infection with phage T7 at a high MOI of 7.

The question arises, why Zorya II only protects against *Autographiviridae* in EHEC, but not against most other tested phage families? Recently, the interaction of ZorA and ZorB of Zorya II with membrane lipids has been analyzed using cell-free transcription-translation in a quartz crystal microbalance with dissipation (TXTL-QCMD), showing that the ZorAB complex localizes at the membrane of the cell poles, whereas the nickase ZorE freely diffuses at the non-pole membrane (46). Consistent with the localization of Zorya II, phage T7 has been proposed to preferentially infect bacterial cells at the poles (46, 47), although the LPS receptor of T7 is widely distributed across the bacterial outer membrane surface (48, 49). T7 phages have very short non-contractile phage tails, and only the first 850 bp segments are initially ejected through the outer membrane (50). Successful ejection of the T7 phage genome into the host cytoplasm requires channel formation involving the phage proteins gp15 and gp16 and the proton motive force (PMF) (50, 51). Thus, both T7 phages and Zorya II rely on an intact PMF for phage DNA translocation and the corresponding phage defense by rotation of the pentameric ZorA tail for ZorE recruitment, possibly explaining why small podoviruses are the preferred targets for Zorya II.

Furthermore, we analyzed the phage spectrum of Druantia III, which is encoded divergently from Zorya II on the OI-172 defense island. Our EOP assays of the Δ*druHE* mutant validated previous findings that Druantia III confers broad-specificity resistance towards many phages (35). In EHEC strain EDL933 Δ*stx1/2*, Druantia III protects against phages of the *Drexlerviridae, Siphoviridae*, *Demerecviridae, Vequintavirinae*, and *Autographiviridae.* However, the protection of Druantia III against most phages was weak, except for the *Epseptimavirus* Bas26 and the *Berlinviruses* Bas67 and Bas68 of the *Autographiviridae*, for which higher EOP differences of 10^2^-10^5^ were observed between the Δ*druHE* mutant and the WT. Previously, high protection levels against >35 susceptible phages were found for heterologously expressed Druantia I and Druantia III in *E. coli* MG1655, as revealed by EOP changes of >10^3^ relative to the MG1655 parent (25, 35). These data indicate that the heterologous expression of Druantia III leads to higher (non-native) levels of phage defense compared to the weak defense conferred by Druantia III in the native EHEC strain. Furthermore, growth data of Δ*druH* and Δ*druE* mutants indicate that both components are required for successful Druantia III-mediated anti-phage defense at the population level.

As a third defense system, we explored the role of the ARMADA II system, which co-localizes with Zorya II and Druantia III in EHEC strain EDL933. Our EOP assays revealed much stronger protection levels by ARMADA II compared to Druantia III against a similar phage spectrum. Specifically, ARMADA II provides strong protection with EOP differences of 10^2^-10^5^-fold between the Δ*armABCD* mutant vs. WT against 26 phages of the *Drexlerviridae, Siphoviridae, Demerecviridae, Vequintavirus* (Bas50), and *Autographiviridae*. Among these are eight phages (Bas5, Bas6, Bas13, Bas29, Bas30, T5, Bas50, Bas67), which are weak targets of Druantia III and very strong targets of ARMADA II. Additionally, ARMADA II showed the highest protection (10^5^ fold-changes) against Bas27, but it does not target Bas26. In contrast, Bas27 is insensitive to Druantia III, whereas Bas26 is highly sensitive. Thus, beyond their similar phage family targets, both systems also act specifically against selected phages, indicating additive activities that results in a broader phage spectrum.

To further study additive and synergistic phage defense, we used EOP and growth assays with AUC calculation of combined mutants deficient in the Zorya II/ Druantia III and Druantia III/ ARMADA II defense pairs, as well as the contribution of the three systems Zorya II/ Druantia III/ ARMADA II to the phage defense of the EHEC strain. The EOP assays revealed weak synergy of the Zorya II/Druantia III combined defenses against nine phages of the *Drexlerviridae, Siphoviridae* and *Vequintaviridae*, but not against *Autographiviridae*, which are strong targets of both individual systems. However, the growth data provide additional evidence for the weak synergy of the Zorya II/ Druantia III defense pair against Bas67 of the *Autographiviridae*. These results are consistent with previous data, which showed that the co-occurring Druantia III and Zorya II systems act synergistically against phages T1, T3, and others (31). However, previous EOP assays were conducted using defense systems, which were heterologously expressed in a laboratory *E. coli* strain (31). In our work, we analyzed the synergistic interactions of the Zorya II/ Druantia III defense pair in the EHEC strain EDL933 Δ*stx1/2* by using deletion mutants, explaining the weaker levels of synergistic interactions compared to the heterologous expression of the systems (31).

Moreover, in the EHEC strain, the ARMADA II system appeared as the dominant defense system, which alone provided the strongest level of protection against a broad spectrum of phages. The EOP assays of the combined Druantia III/ ARMADA II systems showed strong synergistic interactions in the defense against a broad spectrum of 22 phages of different families. Growth assays further validated the additive and synergistic interactions of Druantia III/ ARMADA II defenses against selected phages. Altogether, these data showed that the co-localizing systems Zorya II/ Druantia III and Druantia III/ ARMADA II act to different extents additively or synergistically in the defense against overlapping but not identical phage spectra, together providing more robust immunity against a broader spectrum of phages than alone in the native host. Furthermore, the three systems were also confirmed to be responsible for the strong phage resistance observed in the EHEC strain, since deletion of the systems sensitizes the EHEC strain EDL933 Δ*stx1/2* to the majority of tested phages. The susceptibility of the Δ*zor-dru-arm* mutant towards phage infection is comparable to that of the *E. coli* K-12 strain W3110, indicating that the OI-172 phage defense island might be crucial for the survival of EHEC upon interaction with phages in the ecological context.

The strong synergy between Druantia III and ARMADA II raises the question about the molecular mechanisms of the interaction of both systems in the defense against similar phages. Previously, the molecular basis of the synergistic interactions between Gabija and tmn has been investigated, showing that tmn co-opted the ATPase domain of Gabija for an improved phage defense (31). Similarly, the modular anti-phage systems Druantia III and ARMADA II share effectors with common domains, such as the DExH box NTPases/helicases (DruE and ArmC) with a DUF1998 domain and different C-terminal domains (PLD-like and non-PLD-like nuclease), resembling fused DrmABC components of DISARM systems. Thus, DruE and ArmC could cooperate in the translocation or degradation of the phage DNA for improved phage defense. Additionally or alternatively, the putative helicase/nuclease activities provided by DruE and ArmC might also function in the context of the accessory factors of the respective other system.

Due to their related compositions with respect to domains or subunits, we hypothesize that DruE of Druantia III, ArmC of ARMADA II, and the combined DrmA, DrmB, and DrmC proteins of DISARM systems might function via a similar mode of action during the anti-phage defense. In agreement with the target phages of Druantia III and ARMADA II, the DISARM I and II systems also provide protection against a broad spectrum of phages, including *Siphoviridae, Myoviridae*, and *podoviruses* (22–24). It was proposed that DISARM systems recognize single-stranded overhangs of phage DNA substrates, but not sequence-specific motifs, to circumvent immune evasion mechanisms of phages by escape mutations and to avoid autoimmunity (22). Based on their broad spectra of target phages, we anticipate that DruE and ArmC also recognize the structural context of the DNA substrate, but do not degrade phage DNA in a sequence-specific manner.

However, the three nuclease-based defense systems (Zorya II, Druantia III, and ARMADA II) did not provide immunity against *Tevenviridae* (*Tequatrovirus* and *Mosigvirus*), which are known for their hypermodified phage genomes, including 5-hydroxymethylcytosines (hmCs) and 5-hydroxymethylglucosylcytosines (ghmCs) for *Tequatrovirus* and 5-hydroxymethyl-arabinosylcytosines for *Mosigvirus* (40, 41). The roles of phage DNA modifications for several nuclease-based defense systems have been studied using phage T4 mutants deficient in hmC and ghmC modifications. The PFU assays showed that the presence of phage T4 DNA modifications abolished the Druantia III-mediated defense in vivo (35). Thus, Druantia III and ARMADA II most likely recognize unmodified phage DNA. Our current studies are directed at elucidating the molecular mechanisms of the Druantia III and ARMADA II-based anti-phage defense.

## MATERIALS AND METHODS

### Bioinformatics analysis for the distribution of the OI-172 defense island in *E. coli*.

Public data were investigated concerning the co-occurrence of the genes encoding the three phage defense systems Zorya II, Druantia III, and ARMADA II in pathogenic and non-pathogenic *E. coli* strains. Therefore, all available complete *E. coli* genome sequences (n = 3792) from NCBI RefSeq (https://www.ncbi.nlm.nih.gov/refseq/, accessed on 6 March 2025) (38) were retrieved. The presence of single genes was determined using abricate (https://github.com/tseemann/abricate, with mincov = 50, minid = 15) against a custom database of single defense genes of the gene locus *z5894* to *z5902* from EHEC strain EDL933 (NCBI RefSeq assembly GCF_000006665.1).

Based on the results, only strains comprising at least one complete defense system of the OI-172 defense island (i.e., Zorya II, Druantia III, or ARMADA II) were integrated into further analyses (n = 467, **Table S2**). Strains were characterized using BakCharak (https://gitlab.com/bfr_bioinformatics/bakcharak), which implements the determination of the genoserotype based on the CGE SeroTypeFinder (52) and the EcOH database (53) as well as the MLST typing tool mlst (https://github.com/tseemann/mlst), the ClermontTyping for determination of the phylogroup (https://github.com/A-BN/ClermonTyping) (54), the virulence finder database (55), and AMRFinderPlus v4.0 (56) for determination of the *E. coli* pathotype. Sequence annotation was performed using bakta (57).

Sequence extraction was performed using the extract_genes_ABRicate script (https://github.com/boasvdp/extract_genes_ABRicate) based on the results of an abricate analysis comprising the whole region *z5894* to *z5902* from the EHEC strain EDL933. A distance matrix and the respective UPGMA tree were calculated using mafft (options: --retree 0 –treeout, https://mafft.cbrc.jp/alignment/software/treeout.html). Tree visualization was done using https://microreact.org/. For sequence comparisons of the three defense systems, only 424 strains with all nine genes of the OI-172-associated defense island were used from the first abricate analysis. Two sequences were excluded due to an unusual length of the extracted sequence (GCF_017165395.1) and a duplication in the ARMADA II system (GCF_016942635.1).

### Bacterial strains and cultivation

All bacterial strains, plasmids, phages and primers used in this work are listed in **Tables S6, S7, and S8**. *E. coli* strain DH5α was used as cloning host for plasmid constructions. Additionally, *E. coli* K-12 strains W3110 and MG1655ΔRM were used for EOP assays and for phage propagation (34). The EHEC O157:H7 strain EDL933 Δ*stx1/2* was used as host for construction of phage defense mutants (58) and is referred to as EDL933 wild type (WT) throughout the manuscript, tables, and figures. For growth assays, *E. coli* strains were cultivated at 37°C in LB at an optical density of 600 nm (OD_600_) under vigorous agitation. Statistical analysis was performed using the Student’s unpaired two-tailed *t*-test for two samples of unequal variance, as indicated in the figure legends by *p*-values for each comparison.

### Phage propagation, EOP, and phage infection assays in liquid medium

The 44 phages used in this work for the determination of the phage spectrum and synergy of the three phage defense systems are listed in **Table S7** (34). The phages were propagated in 20 mL LB using the *E. coli* MG1655ΔRM strain lacking all known *E. coli* K-12 restriction systems (34), which was cultivated to an OD_600_ of 0.3 and inoculated with the phage at a MOI of 0.1. Upon phage propagation, the OD_600_ of the *E. coli* MG1655ΔRM culture decreased to 0.1, and 100 µl chloroform was added for 15 min, followed by centrifugation (5000 rpm, 15 min, 4°C) and sterile filtration of the phage lysate. The phage titer was determined using the top agar assay on regular Petri dishes (9.4 mm). The LB top agar contained 0.5% agar and was freshly prepared. About 3 ml of top agar was inoculated with the *E. coli* strain MG1655ΔRM as susceptible host strain, which was mixed with serial dilutions of the phage lysates, and poured as an overlay on solid LB agar plates. After overnight incubation of the LB top agar plates, plaque-forming units (PFUs) were counted for each phage dilution, and the titer of the phage lysate was determined.

Quantitative EOP assays were applied to determine the phage spectrum and synergistic interactions of the defense systems. For EOP assays, larger square Petri dishes (12 cm × 12 cm) were used, and 10 ml of top agar inoculated with 200 μl of an overnight culture of *E. coli* EDL933 Δ*stx1/2* WT or defense system mutant strains to overlay the solid LB agar plates. Afterwards, serial dilutions of phage lysate stocks were prepared in sterile 0.9% NaCl solution, and 5 μl of each dilution was spotted onto the top agar plates and dried. The plates were incubated at 37°C for 24 hours to determine PFUs in the serial dilutions. The EOP of a phage on the defense mutant strain was determined by calculating the ratio of PFUs of the defense mutant versus the EDL933 Δ*stx1/2* (WT) strain. EOP assays for each phage and mutant strain were performed in 3-4 biological replicate experiments.

The synergy or additive protection between the combined systems mutants (Δ*zor-dru*, Δ*dru-arm*, Δ*zor-dru-arm*) versus the sum of the protection of the single system mutants was calculated according to the formula: [log_10_EOP (WT/combined mutant)] – [(log_10_EOP (WT/single mutant1)) + (log_10_EOP(WT/single mutant2))], which indicates the synergy score. Synergistic interaction is given when [log_10_EOP (WT/combined mutant)] > [(log_10_EOP (WT/single mutant1)) + (log_10_EOP (WT/single mutant2))]. Additive protection is given when [log_10_EOP (WT/combined mutant)] < [(log_10_EOP (WT/single mutant1)) + (log_10_EOP(WT/single mutant2))]. The synergistic or additive interaction of the defense systems is shown as a yellow-green color code in **Fig. 5**, and the synergy values are presented in **Table S4**.

For liquid phage infection experiments at the population level, the EHEC strains were grown in LB to an OD_600_ of 0.08 and infected with different MOIs of the phages to monitor the growth and collapse of the bacterial populations. The average growth curves from 3-4 biological replicates of non-infected controls and infected cultures were used for the calculation of the AUC for quantitative comparison of the synergy of the combined versus single system mutants. The formula ((OD1+OD2)/2)*(t2-t1) was applied for the determination of the AUC for each time point of OD_600_ measurement along the growth, which was then summed up for the total AUC of each growth curve. The total AUC of the average growth curve of the phage-infected strain EDL933 Δ*stx1/2* WT or defense mutant strains was normalized to the AUC of the average growth curve of the non-infected control for each strain, which was set to 100%.

### Construction of the phage defense mutants and complemented strains in EHEC strain EDL933 Δ*stx1/2*

The single mutants and multiple defense system deletion mutants of the Zorya II, Druantia III, and ARMADA II systems located on OI-172 were constructed as clean (scarless) deletions in the EHEC strain EDL933 Δ*stx1/2* by using the one-step inactivation method of Datsenko and Wanner (59). First, the chloramphenicol resistance (*cat*) cassette, which is flanked by FRT (FLP recognition target) sites, was amplified by PCR from the pKD3 plasmid using gene-specific primers **(Table S8)** containing ∼50 bp homology arms corresponding to the sequences immediately up- and downstream of the target gene in the EDL933 Δ*stx1/2* genome. Next, the single or multiple defense system genes were replaced by the *cat* cassette via the pKD46-encoded λ red recombinase.

For mutant constructions, purified PCR products of the *cat* cassette (1 µg) were electroporated into electrocompetent *E. coli* EDL933 Δ*stx1/2* pKD46 cells using a 0.1 cm gap cuvette at 1.8 kV. Following electroporation, cells were recovered in 1 mL LB medium at 37 °C for 90 minutes and plated on LB agar supplemented with chloramphenicol (25 µg/mL). Plates were incubated overnight at 37°C. Putative mutants were screened by colony PCR using primers flanking the integration site **(Table S8)**. The pKD46 plasmid was subsequently cured by incubation at 42°C, and scarless mutants were obtained via FLP-mediated excision of the *cat* cassette using the pCP20 plasmid, as previously described (59).

For complementation of the EDL933 Δ*stx1/2* Δ*druE*, Δ*armC*, and Δ*zorE* deletion mutants, the low copy number plasmid pWKS30 was used (60), and the *druE*, *armC*, and *zorE* genes were cloned into pWKS30 by Gibson assembly (61). Thereby *armC* and *zorE* were amplified from the EDL933 Δ*stx1/2* genome. The vector pFL-*druE* was used as a template for the amplification of *druE*. For the construction of the pFL-*druE* vector, synthetic gene blocks encoding *druE,* codon-optimized for insect cell expression, were stepwise assembled and integrated into the pFL vector again using Gibson assembly (61). The pWKS30 plasmids encoding *druE*, *armC*, or *zorE* were electroporated into the respective defense mutants. The *druE*, *armC*, and *zorE* complemented strains were selected on LB agar plates containing ampicillin and approved by growth assays with selected phages to restore phage defense after induction by 200 µM IPTG as compared to the respective mutants.

## ACKNOWLEDGEMENTS

This work was supported by a grant from the Deutsche Forschungsgemeinschaft (AN746/8-1) and overheads of the ERC CoG grant (GA 615585) to H.A. We are grateful to Alexander Harms and Thomas Gaudin for providing the phages for this work. We thank Ulrich Dobrindt for construction of the initial *druE, druH*, and *druHE* mutants and for the comments to the manuscript. We are thankful to the bachelor students Leonie Nowara, Sinem Ece Ersöz, Nermin Ibrahim, and Timon Sturm for their help in growth assays of the phage defense mutants. We thank Franziska Kiele for the excellent technical assistance.

## DATA AVAILABILITY STATEMENT

The authors confirm that the data supporting the findings of this study are available within the article and its supplementary materials.

## Supplemental Figure legends

**Fig. S1A-G. Growth curves of combined and single defense system mutants deficient in the Zorya II, Druantia III, and/or ARMADA II systems**. The growth curves of the Δ*zorABE,* Δ*druHE*, and Δ*armABCD* single system mutants and of the Δ*zor-dru*, Δ*dru-arm*, and Δ*zor-dru-arm* combined system mutants were monitored in LB after infection with the phages Bas6 (A), Bas20 (B), Bas26 (C), Bas27 (D), Bas30 (E), Bas 50 (F), and Bas 67 (G) at different MOIs. Average values of three biological replicates of the growth curves for the defense mutants are presented for the nine phages with error bars representing the SD. The statistics were calculated for the infected defense mutants versus the uninfected control using the two-tailed unpaired t-test. *P* values: *, *p* < 0.05; **, *p* < 0.01; ***, *p* < 0.001. The growth curves were used for AUC calculation during the five hours of growth as presented in **Fig. 6A-C**.

## Supplemental Table legends

**Table S1. Distribution of the co-localizing defense systems Zorya II, Druantia III, and ARMADA II in *E. coli* and other proteobacteria based on the KEGG database**. OI-172 of the EHEC strain EDL933 encodes the three defense systems Zorya II (*zorABE*), Druantia III (*druHE*), and ARMADA II (*armABCD*), which co-localize in STEC EHEC O157:H7, O55:H7, and O145:H28 strains as well as in the non-pathogenic *E. coli* strains ATCC8739 and SMS-3-5. Additionally, the three defense systems are incomplete in several other proteobacteria, which were identified using the KEGG database (https://www.kegg.jp/kegg/) upon searching for *druE* homologs of *E. coli* strain EDL933 using the SSDB Gene Cluster Search (https://www.kegg.jp/ssdb-bin/ssdb_gclust?org_gene=ece:Z5898).

**Table S2. Distribution of the co-localizing defense systems Zorya II, Druantia III, and ARMADA II in 467 genomes of pathogenic and non-pathogenic *E. coli* from the NCBI RefSeq collection**. The three defense systems Zorya II (*zorABE*), Druantia III (*druHE*), and ARMADA II (*armABCD*) are encoded on OI-172 of the EHEC strain EDL933. The sequences of the OI-172-associated defense systems of EDL933 were compared with the complete genome sequences of 3792 *E. coli* strains retrieved from the NCBI RefSeq database. The % sequence identities of the nine defense genes of EDL933 to the identified defense genes of the *E. coli* genomes are presented. For some genes, 2 values are given as % sequence identity, indicating that a second homolog of the defense gene is present in the same genome. Only genome sequences are shown with sequence similarity to at least 1, 2, or 3 complete defense systems of OI-172 (e.g., Zorya II, Druantia III, ARMADA II) of strain EDL933. The *E. coli* strains harboring the defense systems were classified according to their MLST sequence type (ST), O- and H-antigen serotypes, Clermont type phylogroups, and pathotype as derived from the BakCharak analyses. Numbers for the defense genes *zorE, zorB, zorA, druH druE, armA, armB, armC*, and *armD* represent the results from the abricate analysis (coverage) using EHEC strain EDL933 (GCF_000006665.1) as reference. Abbreviations: ONT = O not typeable, aEPEC = atypical enteropathogenic *E. coli*, STEC = Shiga toxin-producing *E. coli*, noDEC = no diarrheagenic *E. coli*.

**Table S3. EHEC strain EDL933 Δ*stx1/2* is more resistant to most phages of the BASEL collection compared to the *E. coli* K-12 strain W3110, as revealed by PFU/EOP assays**. The log_10_ EOP values were determined in PFU assays as fold-changes of the phage dilutions of plaque formation of the EDL933 Δ*stx1/2* versus the W3110 strain against a panel of 44 phages of the BASEL collection as described in Materials and Methods. The PFU experiments were performed in 3-4 biological replicates for the strains W3110 and EDL933 Δ*stx1/2*.

**Table S4. Phage spectrum and synergy of the Zorya II, Druantia III, and ARMADA II anti-phage systems in the EHEC strain EDL933 Δ*stx1/2***. To determine the phage spectrum targeted by the Zorya II, Druantia III, and ARMADA II defense systems, the log_10_ EOP values were determined in PFU assays as fold-changes of the phage dilutions for plaque formation in the strain EDL933 Δ*stx1/2* (WT) versus the *druHE, zorABE*, and *armABCD* mutants against a panel of 44 phages of the BASEL collection as described in Materials and Methods. The EOP differences are indicated by a white-red color code based on the level of protection. The synergy or additive protection of the combined defense systems was analyzed in PFU/EOP assays of the *zor-dru*, *dru-arm*, and *zor-dru-arm* mutants versus the WT. Synergy of anti-phage defense of combined defense systems was defined when EOP values of the combined mutants were above the additive EOP values of the individual defense mutants. The synergy score is given by a yellow-green color code. All PFU experiments were performed in 4 biological replicates for each EDL933 Δ*stx1/2* defense mutant strain. The data are presented as a heatmap in **Fig. 3**.

**Table S5. Complementation of the helicase/nuclease effector mutants of the Zorya II, Druantia III, and ARMADA II defense systems restored their activity in the phage defense.** We analyzed the functionality in the anti-phage defense for the *druE, zorE*, and *armC* helicase/nuclease effector mutants after complementation with pWKS30-encoded *druE*, *zorE*, and *armC*. Specifically, the log_10_ EOP values were determined in PFU assays as fold-changes of the phage dilutions for plaque formation of strain EDL933 Δ*stx1/2* (WT) and the *druE*, *zorE*, and *armC* complemented strains versus the *druE, zorE*, and *armC* mutants, respectively, against a panel of 44 phages of the BASEL collection as described in Materials and Methods. The EOP differences are indicated by a white-red color code based on the level of protection. All EOP experiments were performed in 4-5 biological replicates for each strain. The data are presented as a heatmap in **Fig.3**.

**Table S6. Bacterial strains and plasmids Table S7. Phages used in this study**

**Table S8. Oligonucleotide (primer) sequences**

